# Mast seeding promotes evolution of scatter-hoarding

**DOI:** 10.1101/698761

**Authors:** Rafał Zwolak, Dale Clement, Andrew Sih, Sebastian J. Schreiber

## Abstract

Many plant species worldwide are dispersed by scatterhoarding granivores: animals that hide seeds in numerous, small caches for future consumption. Yet, the evolution of scatterhoarding is difficult to explain because undefended caches are at high risk of pilferage. Previous models have attempted to solve this problem by giving cache owners large advantages in cache recovery, by kin selection, or by introducing reciprocal pilferage of “shared” seed resources. However, the role of environmental variability has been so far overlooked in this context. One important form of such variability is masting, which is displayed by many plant species dispersed by scatterhoarders. We use a mathematical model to investigate the influence of masting on the evolution of scatter-hoarding. The model accounts for periodically varying annual seed fall, caching and pilfering behavior, and the demography of scatterhoarders. The parameter values are based mostly on research on European beech (*Fagus sylvatica*) and yellow-necked mice (*Apodemus flavicollis*). Starvation of scatterhoarders between mast years decreases the population density that enters masting events, which leads to reduced seed pilferage. Satiation of scatterhoarders during mast events lowers the reproductive cost of caching (i.e. the cost of caching for the future rather than using seeds for current reproduction). These reductions promote the evolution of scatter-hoarding behavior especially when interannual variation in seed fall and the period between masting events are large.

## Introduction

Masting, or periodic, synchronized production of abundant seed crops, is a common reproductive strategy of many plants (Kelly 1994, Kelly and Sork 2002) and a classic example of a pulsed resource (Ostfeld and Keesing 2000, Yang et al. 2010). Masting provides foundational, yet unstable resource levels, creating cycles of feast and famine in food webs (Clark et al. 2019). These cycles strongly influence behavior and life-history strategies of consumers. For example, animals migrate to track populations of masting plants (Jenni 1987), enter diapause to survive lean periods between mast-events (Maeto and Ozaki 2003), and increase reproduction in anticipation of masting (Boutin et al. 2006, Bergeron et al. 2011). However, despite decades of research, many impacts of masting on consumers remain poorly understood (Clark et al. 2019). Here we use a mathematical model to show that masting can play an important but overlooked role in the evolution of a widespread animal behavior: scatter-hoarding.

Scatter-hoarding is defined as caching seeds for future consumption in many small, widely-dispersed caches (Vander Wall 1990). This caching strategy is used by numerous species of animals, most notably by rodents and corvids (Pesendorfer et al. 2016, Lichti et al. 2017, Gómez et al. 2019). Scatterhoarders provide essential seed dispersal services in many ecosystems throughout the world. According to a recent review, there are 1279 species of plants known to rely on this mode of seed dispersal, although this number is certainly underestimated (Gómez et al. 2019). However, even though scatter-hoarding is so widespread, the evolutionary advantage of this behavior is not obvious because the caches are undefended and often suffer very high rates of pilferage (Schmidt and Ostfeld 2008; Jansen et al. 2012; Zwolak et al. 2016; Dittel et al. 2017). Thus, scatter-hoarding appears vulnerable to cheating by non-caching pilferers (Andersson and Krebs 1978; Smulders 1998; Vander Wall and Jenkins 2003).

First attempts to solve this problem focused on the role of the owner’s advantage. According to a model by Andersson and Krebs (1978), scatter-hoarding can evolve when cache owners are substantially more likely to recover caches than are naive foragers. Empirical estimates of the owner advantage vary widely but appear relatively high in scatter-hoarding birds (particularly those that rely on specialized spatial memory: e.g. Brodin 2010) and quite low in mammals. In most studies on rodents, cache owners are only 2-4 times more likely to recover their caches when compared with naïve individuals (Jacobs and Liman 1991; Jacobs 1992; Briggs and Vander Wall 2004; Thayer and Vander Wall 2005; Vander Wall et al. 2006; 2008; Hirsch et al. 2013; Gu et al. 2017; see also Steele et al. 2011). In many systems, the owner’s advantage appears to be insufficient to prevent substantial cache loss to pilferage (2-30% lost per day, according to a review by Vander Wall and Jenkins 2003, though this rate might vary depending on environmental characteristics like soil moisture: Vander Wall 2000).

A later model by Vander Wall and Jenkins (2003) suggested that caching can represent an adaptive, stable strategy when all caches are reciprocally pilfered by scatter-hoarding animals with overlapping home ranges (see also Smulders 1998). Under this scenario, caches represent a collective resource used by selfish individuals (Vander Wall and Jenkins 2003). The reciprocal pilferage hypothesis predicts that animals are unlikely to avoid pilferage, but can compensate for it by pilfering caches of other individuals. As a corollary, individuals should invest in their pilfering tactics rather than in theft-reducing strategies (but see e.g. Dally et al. 2006, Steele et al. 2008, Galvez et al. 2009, Shaw and Clayton 2013, Hirsch et al. 2012, Muñoz and Bonal 2011 for examples of potentially costly behaviors aimed to reduce pilferage).

Environmental variability represents an additional, potent mechanism for scatter-hoarding that has been largely overlooked in the existing models. Such variability is pervasive in ecosystems dominated by plants that produce scatterhoarder-dispersed fruits because such plants usually show pronounced masting (Herrera et al. 1998; Vander Wall 2001). Examples of scatterhoarder-dispersed masting plants can be found in the tropics (Norden et al. 2007; Mendoza et al. 2018), deserts (Meyer and Pendleton 2015; Auger et al. 2016), and in temperate zones (Koenig and Knops 2000; Schauber et al. 2002; Shibata et al. 2002). While studies of masting have often emphasized the benefit of masting to plants in terms of reduced per capita seed predation (“predator satiation”: Kelly 1994), masting also has important effects on consumer population dynamics that can feedback to affect the evolution of caching. In particular, the cycles of satiation and starvation induce striking fluctuations in consumer population size (Ostfeld and Keesing 2000; Yang et al. 2010; Bogdziewicz et al. 2016). Typically, masting triggers a temporary increase in consumer population size followed by a pronounced crash. Thus, when the next mast year comes, seed to consumer ratios are particularly high (Kelly 1994; Ostfeld and Keesing 2000).

We use a mathematical model to investigate the influence of mast-related fluctuations in scatterhoarder population size on the evolution of scatter-hoarding. The model mimics interactions between a masting tree and a scatterhoarding rodent. The scatterhoarders consume or cache harvested seeds and pilfer or recover their own caches over years that differ in both the magnitude of seed fall and the number of competing consumers. Previous models demonstrated that caching is influenced by the owner’s advantage in cache recovery and the probability that scatterhoarders survive long enough to use the caches (Andersson and Krebs 1978; Smulders 1998; Vander Wall and Jenkins 2003). However, both the proportion of recovered seeds and scatterhoarder survival depend on the magnitude of seed fall and the resulting fluctuations in population size (Pucek et al. 1993, Theimer 2005, Zwolak et al. 2016). Thus, we expand on previous models by including the effects of environmental variability resulting from mast seeding on caching behavior. We do this by treating the proportion of seeds that are cached, rather than immediately consumed, as an evolving trait and examining how the evolutionarily stable strategy of this caching behavior varies with (1) masting intensity, (2) the frequency of mast years, (3) the owner’s advantage in cache recovery, and (4) the survival of scatterhoarders. Our results demonstrate that mast-related fluctuations in scatterhoarder population size reduce both the risk of cache loss to pilferers and the reproductive cost of caching (i.e. the cost of caching seeds for future use rather than using seeds for current reproduction), thus promoting the evolution of scatter-hoarding.

## Methods

### Modeling approach

We consider a population of scatterhoarders that experience three distinct periods of seed availability in each year: fall, winter/spring, and summer. During the fall, seeds become available, and scatterhoarders gather and either immediately consume or cache them. Energy from consumed seeds contributes to reproduction while cached seeds may be recovered for use during the subsequent winter/spring. During the winter/spring, scatterhoarders are sustained by seeds from their own caches or seeds they pilfer from the caches of other scatterhoarders. As caching behavior may not always be favored, we implicitly assume the availability of other winter resources that prevent population extirpation. During the summer, scatterhoarders survive and reproduce using resources other than seeds. Caching behavior for an individual is represented by the threshold *T*, such that the individual consumes up to *T* seeds during the fall and caches the rest. We determine the conditions under which caching behavior is favored by solving for the evolutionary stable strategy (ESS) of *T*. The central conundrum is that the strategy of scatter-hoarding appears vulnerable to cheating by non-caching pilferers, who may invade the population and outcompete caching individuals. If the population is monomorphic for a particular caching threshold, then this threshold value is an ESS if this population cannot be invaded (outcompeted) by individuals with any other threshold value.

### Model description

During fall (period 1), there is seed fall of *S*(*t*) from the primary seed source. Seeds are gathered at a rate proportional to the density *n*_*1*_(*t*) of scatterhoarders during this period. The proportionality constant *a*_*1*_corresponds to the per-capita (i.e., per scatterhoarder) seed harvest rate. Seeds are also lost to other sources (e.g. competitors, germination, decay, etc.) at a per-capita rate of *L*_*1*_. If all seeds are gathered or lost to other sources by the end of the fall, then the amount of seed gathered per individual equals:

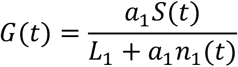

All seeds above a threshold, *T*, are cached by individuals for later in the year. The seeds which are not cached, min{*G(t),T*}, are used for survival and reproduction. The number of offspring produced by an individual, *R*_1_(*t*), at the end of the fall is a saturating function of min{*G(t),T*}, with a maximal number of offspring *b* and a half saturation constant *h* (i.e. *h* is the amount of resources required to produce *b*/2 offspring). The fraction of adults surviving from the first period (fall) to the second period of the year (winter/spring) equals *s*_1_. Thus, the total density *n*_2_(*t*) of individuals entering winter/spring equals

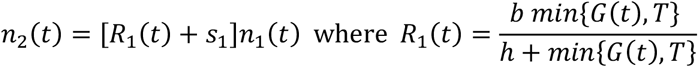

The main resource available to individuals during winter/spring is the total size of seed caches max{*G*(*t*)-*T*,0}*n*_1_(*t*). Owners of the cached seed who survived gather their cached seed at a rate proportional to the size of their seed caches. This proportionality constant *a*_2_corresponds to the per-capita rediscovery and use rate of their caches. All other individuals are assumed to pilfer seed from others’ caches at a per-capita rate *a*_pil_. Seeds are lost to other sources at a per-capita rate of *L*_2_. If all cached seeds are gathered or lost by the end of winter/spring, then the fraction of caches recovered by its owner given that the owner survived from fall to winter/spring is

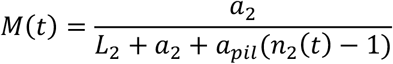

while the fraction of caches that was pilfered by each non-owner is

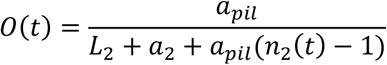

For seed caches whose owner died, the fraction that was recovered by a living non-owner is

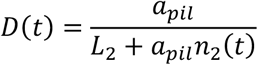

The total amount of cached seed gathered by a surviving individual from fall is the amount of seed recovered from its own caches plus the amount of seeds pilfered from the caches of other surviving individuals and the caches of deceased individuals:

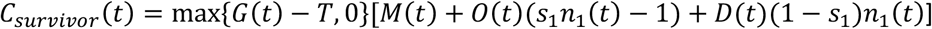

Since individuals that were born at the end of the fall had no opportunity to cache, the total amount of cached seed gathered by these individuals in the spring/winter is only the amount pilfered from the caches of either surviving or deceased individuals:

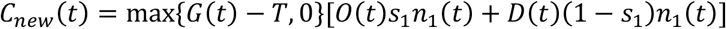

If *s*_*2*_is the fraction of individuals surviving to summer (period 3), then the density of individuals entering summer equals

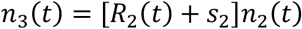

where *R*_2_(*t*) is the population-level per-capita fecundity corresponding to the weighted combination of reproductive contributions of individuals surviving from fall and new individuals born at the end of fall (for simplicity, we assume the same value of *h* for surviving and new born individuals):

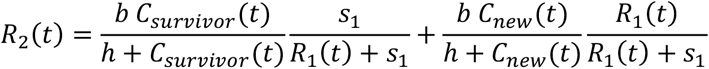

During this final period of the year (summer), individuals rely on other resources with availability *A* to reproduce and survive with probability *s*_*3*_ (these resources represent alternative foods, such as fungi, invertebrates, or seeds of other plant species; adjusted to obtain population dynamics consistent with patterns observed in the field). Thus, the density of individuals entering the fall of the next year equals

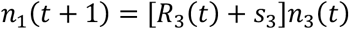

where *R*_*3*_(*t*) is the per-capita reproduction. We model this per-capita reproduction using a Beverton-Holt function

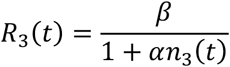

where *β* is the maximal summer fecundity and *α* determines the strength of intraspecific competition. By composing the equations across the three periods of the year, the yearly update rule for population densities at the beginning of fall is

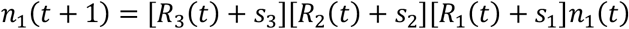

We modeled seed fall S(t) in the fall as a periodic function of time where the period P corresponds to the time between masting years. In the masting years, *S(t)=S*_*high*._, next year *S(t)=S*_*min*_ (typically, seed crops produced after mast years are particularly scant: Pearse et al. 2016, Bogdziewicz et al. 2020), then *S(t) = S*_*low*_ until another mast year. Our analysis assumes that the average seed output, (*S(1)+S(2)+…+S(P))/P*, is fixed and what varies is the proportion of total seed output in the masting year. Higher intensity of masting means more seeds during the masting year, but concomitantly fewer seeds in other years (as opposed to just increasing seed output in masting years with no effect on seed production in other years). Similarly, when we vary the number of years between masting events, the average seed output remains the same (i.e., longer intermast interval corresponds to higher seed production in mast years).

Figure 1 illustrates typical dynamics of the model for the baseline parameters described below. In this figure, seed masting occurs every four years (Fig. 1a) and leads to a stable, four-year population cycle (Fig. 1b). Population densities (Fig. 1b) exhibit seasonal as well as yearly variation. Highest densities are reached at the end of the masting year (year 1) and crash to low densities the ensuring years (years 2-4). For lower caching thresholds, caching occurs in all years except the year after a masting event (green bars in Fig. 1a). Table A1 in Appendix A lists all parameters for the model and their meaning.

**Fig. 1.**
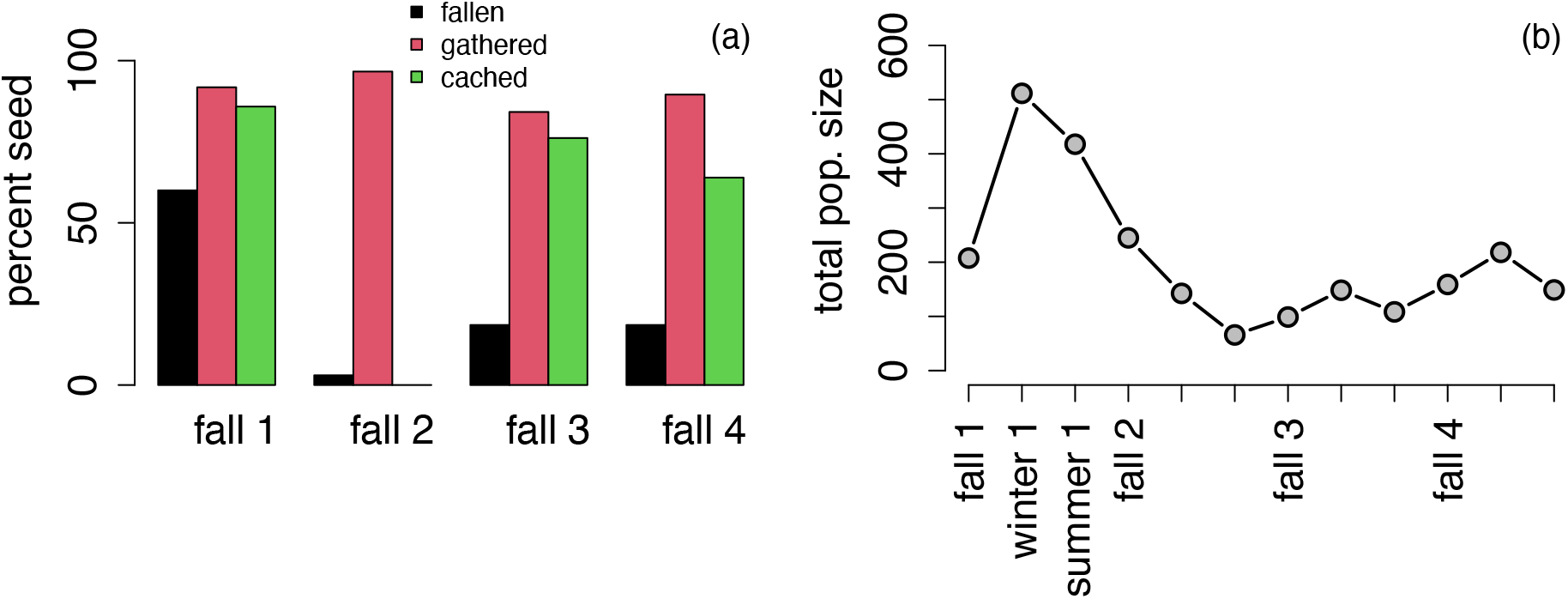
The annual dynamics of fall seeds (a) and seasonal population dynamics (b) when masting years occur every 4 years. Both (a) and (b) correspond to a stable, periodic solution of the model. In (a), the percent of seeds that fall each year from all of the seeds in 4 year masting interval year are plotted as black bars; the first year corresponds to the masting year that is followed by years of lower seed fall. The percent of fallen seed that are gathered each year correspond to the red bars, while the percent of gathered seed that is cached are the green bars. In (b), the total population densities vary intra- and inter-annually; the highest densities occur in the winter/spring period of the masting year after which the population densities crash to lower densities. Parameter values as described in the main text with a 6-fold owner’s advantage in cache recovery.

### Model Parameters

The parameter values are based mostly on research on European beech (*Fagus sylvatica*) and yellow-necked mice (*Apodemus flavicollis*). *Apodemus* mice are among the most important seed predators and scatterhoarders in Eurasia (e.g. Muñoz and Bonal 2011, Shimada et al. 2015, Yang et al. 2019, Wróbel and Zwolak 2019). While we found it useful to base our parameter estimates on a specific, reasonably well-studied system, we also performed a global sensitivity analysis for our main conclusions (Appendix B). Specifically, for each of main results, we reran the simulations 100 times with each parameter, call it *x*, chosen independently from a uniform distribution on the interval [x/1.5,1.5x].

The parameters *a*_1_ and *L*_1_ (per-capita harvest rate and per-capita seed loss) may be reduced to the single parameter *L*_1_/*a*_1_. Rearranging the equation for the amount of seeds gathered *G* yields 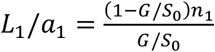. The parameters *n*_1_ and *n*_2_ were taken to be the average species-wide density for A*podemus flavicollis*: 17.7 individuals/ha (Jones et al. 2009). Estimates of the proportion of seeds removed from the forest floor (*G*/*S*_5_) tend to be variable with Zwolak et al. (2016) reporting 78% seed removal during mast years and 91% seed removal during non-mast years for *A. flavicollis*, and Le Louarn and Schmitt (1972) reporting 61% and 74% seed removal by *Apodemus sylvaticus* during two different years. We selected the average value of 76% as our estimate of seed removal. Thus, *L*_1_/*a*_1_ = 5.59. In all cases where *L*_1_ and *a*_1_ were treated as separate parameters, we used *a*_1_ = 1 and set *L*_1_ equal to our choice for *L*_1_/*a*_1_.

The parameters *a*_*pil*_ and *L*_2_ may similarly be reduced to *L*_2_/*a*_*pil*_. Zwolak et al. (2016) estimated the recovery of seeds from artificial caches to be 54% during nonmast years and 5% during mast years, which we assumed to be roughly equivalent to the proportion of seeds recovered from an abandoned cache (*D*/ max{*G*(*t*) − *T*, 0}). We then estimated *L*_2_/*a*_*pil*_ to be between 15.1 and 336.3. In our analysis, we set this value to the upper end of this range (300) as this leads to more conservative estimates of when caching evolves. In all cases where *L*_2_and *a*_*pil*_ were treated as separate parameters, we used *a*_*pil*_ = 1 and set *L*_2_ equal to our choice for *L*_2_/*a*_*pil*_.

We let *a*_2_ = 3 (when *a*_*pil*_ = 1). This value approximates the results of several studies on scatterhoarding rodents (Vander Wall et al. 2006, 2008; Thayer and Vander Wall 2005; Hirsch et al. 2013) that documented seed removal rates by cache owners and naïve foragers. However, we explored scenarios with both higher and lower cache owner’s advantage (see Results).

We assumed a maximum litter size of 11 individuals (Macdonald and Tattersall 2001), with one breeding event per period (2-3 litters per year: Pucek 1984). Assuming that half the population are female and half of the individuals born are female, this yields *b* = 5.5.

Half-saturation constant for mid-year reproduction (*h*) was set as 124 seeds/offspring multiplied by half the maximum number of female offspring (*b*). This value was calculated on the basis of energy contents of beech seeds (Grodziński and Sawicka-Kapusta 1970), energy requirements of yellow-necked mice (0.60 kcal/g/day: Jensen 1982; average body mass of yellow-necked mice is 28.3 g: AnAge), and typical costs of reproduction-related energy expenditure in small mammals (25% increase in energy expenditure during gestation and 200% increase during lactation: Millar 1978; 1979; Gittleman and Thompson 1988; Sikes 1995; Zhu et al. 2015), given the length of gestation and lactation in yellow-necked mice (26 and 22 days, respectively: AnAge). Note that the link between food availability and reproduction limits winter breeding to masting events (which are known to result in winter reproduction in our and related study systems: Jensen 1982, Pucek et al. 1993, Wolff 1996, Ostfeld et al. 1996).

We used 77.5% as the yearlong monthly survival rate (calculated from data on winter survival in Pucek et al. 1993: see also Jensen 1982 for similar values). We assumed that each period lasts four months, yielding 0.775^4^=36.1% as the survival rate for each period (*s*_1_, *s*_2_, and *s*_3_). Winter survival rates in Pucek et al. (1993) are similar to monthly summer survival rates reported or calculated from other studies (e.g. Bujalska and Grüm 2008, Sozio and Mortelliti 2016), thus we assumed equal survival across all seasons in our initial scenario, but examined how relaxing this assumption affects caching rates (see Results).

In principle, food availability affects both reproduction and survival in a manner that depends on life history allocation. That is, an organism can allocate most of its energy budget to enhanced survival or enhanced reproduction, or a blend of the two. Predicting this life history allocation is complex (Roff 2002), thus rather than attempt to predict the optimal allocation (which should depend on optimal caching and vice versa), we draw on the natural history of the system to argue that as a first pass, it is more important to examine how food consumption affects fecundity as opposed to survival. An increase in food consumption clearly increases fecundity. In contrast, we assume that the consumer has alternative food sources (see above) that are sufficient to allow it to survive adequately even if it does not allocate any additional energy from the focal seed source towards survival. We further assume that allocating extra energy to increased survival is not very effective (in our system) because survival also depends heavily on predation, disease, etc. (Jędrzejewska & Jędrzejewski 1998). In this scenario, survival depends little on the amount of the focal seed source consumed. This is also in line with numerous empirical studies reporting that rodents allocate extra energy to increase reproductive output rather than survival (meta-analyzed by Prevedello et al. 2013). In addition, a critical point for relating food to demography is that in short-lived, fecund animals such as rodent scatterhoarders elasticity for survival is low whereas elasticity for reproduction is higher (Heppell et al. 2000), which means that even if survival does vary, this variation does not affect population growth as much as do changes in reproduction. Accordingly, we focus on effects of food on reproduction, and simplify the analysis by assuming that survival is a parameter that is constant across years. However, we do vary survival across all years (see Results).

### Numerical Methods

To identify the evolutionary stable caching strategies, we examined whether a small mutant subpopulation using the caching threshold *T*_*m*_ can invade a resident population using the caching threshold *T*. When the mutant subpopulation densities *m*_*i*_*(t)* in each of the periods *i=1,2,3* are sufficiently small, the effect of the mutant population on the resident population and itself is negligible. Hence, the dynamics of the mutant in the initial phase of invasion can be approximated by the mutant’s growth rate when the population is composed entirely of residents. We now describe these dynamics.

As the mutant and resident individuals only differ in their caching strategy, the amount of seeds gathered in year *t* by a mutant individual equals the amount of seeds gathered *G(t)* by a resident individual. As for the resident dynamics, yearly update of the mutant’s fall density is of the form

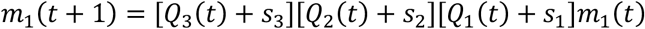

*Q*_1_(*t*) corresponds to the number of offspring produced by a mutant individual during the fall and only differs from the resident in its threshold *T*_*m*_

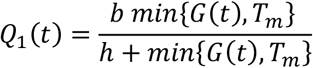

*Q*_2_(*t*) corresponds to the number of offspring produced by a mutant individual during the winter/spring given by a weighted combination due to the fraction of individuals that survived from the fall and individuals born in the fall:

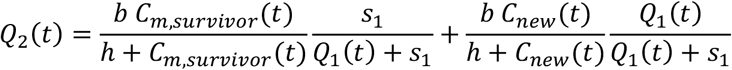

where *Q*_2_(*t*) differs from *R*_2_(*t*) only in its first term due to surviving individuals with the mutant caching strategy: *C*_*m*.*survivor*_(*t*) = max{*G*(*t*) – *T*_*m*_, 0}*M*(*t*) + max{*G*(*t*) − *T*, 0}[*O*(*t*)(*s*_1_*n*_1_(*t*) − 1) + *D*(*t*)(1 − *s*_1_)*n*_1_(*t*)]. Finally, the number of offspring produced by a mutant over the summer is the same as the resident i.e. *Q*_3_(*t*) = *R*_3_(*t*).

Whether the mutants playing strategy *T*_*m*_ are able to invade the residents playing the strategy *T* or not depends on their long-term per-capita growth rate

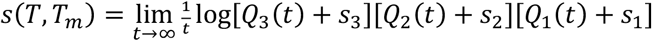

provided the limit exists. Over the parameter space (see previous section) that we simulated, the population dynamics always converged to a periodic solution whose period *kP* is a multiple *k* of the seed masting period *P*. Typically, this multiple was 1 or 2 or 4, the latter two corresponding to period-doubling bifurcations. We developed R code to efficiently approximate these periodic solutions. For these periodic solutions of the resident dynamics, the long-term per-capita growth rate of mutant strategy *T*_*m*_ against resident strategy *T* equals

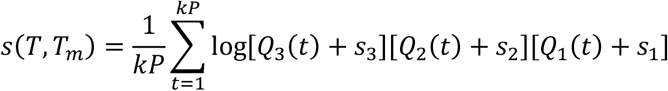

A strategy T is an evolutionarily stable strategy (ESS) for caching if *s*(*T,T*_*m*_)<0 for all strategies *T*_*m*_ ≠T. To find ESSs for caching, we derive in Appendix C an explicit expression for the fitness gradient 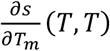 when the resident population is playing threshold strategy *T*. When the fitness gradient is positive, mutants with a higher threshold strategy than the residents can invade while mutants with a lower threshold strategy fail (Geritz et al. 1997). When the fitness gradient is negative, the opposite occurs. As mutants with larger or smaller thresholds fail when invading a resident population playing the ESS, the fitness gradient equals zero at an ESS. Hence, we identified ESSs by iteratively solving for thresholds T at which the fitness gradient 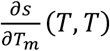 is zero (Fig. C1 in Appendix C).

Our results focus on the fraction of seeds cached (*F*) rather than the caching threshold (*T*), as this quantity is easier to interpret. The relationship between these two measures of caching is given by 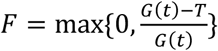. As the amount of seeds gathered G(t) varies from year to year, the percentage of seeds cached when playing the ESS also varies from year to year. We also examine pilferage risk and the marginal reproductive cost of caching. Pilferage risk is the probability (expressed as a percent) that a seed is pilfered from a surviving individual’s cache during winter/spring and equals 100(*n*_2_(*t*) − 1)*O*(*t*). If *F*_*m*_ denotes the percentage of seeds cached by mutant individuals, then the marginal reproductive cost of caching equals the infinitesimal reduction in reproductive output for a mutant individual caching an infinitesimal amount of seeds rather than consuming them i.e., 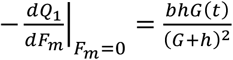.

## Results

Increasing intensity of masting results in decreased fall scatterhoarder population density (i.e., the density that enters masting events; Fig. 2a). This occurs because reproduction is a saturating function of seeds gathered and the reproductive gains of higher seed availability during masting years are outweighed by the reproductive losses due to lower seed availability during non-mast years.

**Fig. 2.**
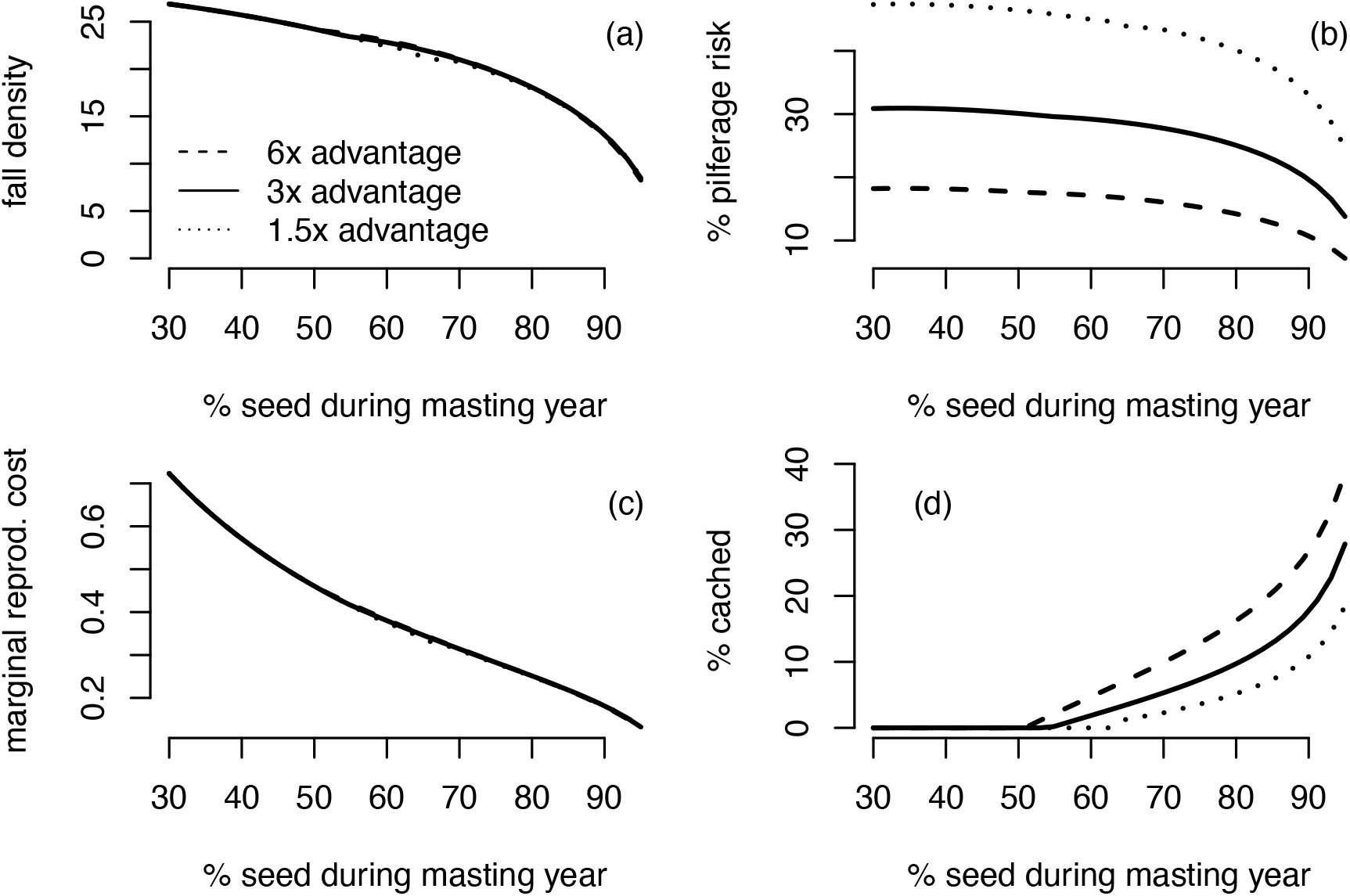
Mast year fall population density in individuals/ha (a), pilferage risk, defined as the probability that cached seed would be pilfered (b), marginal reproductive costs of caching (c), and proportion of seeds cached rather than eaten (d) as a function of masting intensity (expressed as the percentage of total seed production that occurs during mast years), with mast occurring every 4^th^ year. Dashed, solid, and dotted lines represent the magnitude of owner’s advantage in cache recovery (owners 6, 3, and 1.5 times more likely to discover their own caches relatively to naïve foragers). All dependent variables are given at the evolutionary stable caching strategy and its associated periodic population dynamics.

Increasing masting intensity also reduces the risk that a cached seed would be pilfered (Fig. 2b), particularly when a high proportion of seeds are produced during mast years. The responses of pilferage risk and fall density are correlated (see, also, Figs. 3 and 4) because lower population density means fewer pilferers.

**Fig. 3.**
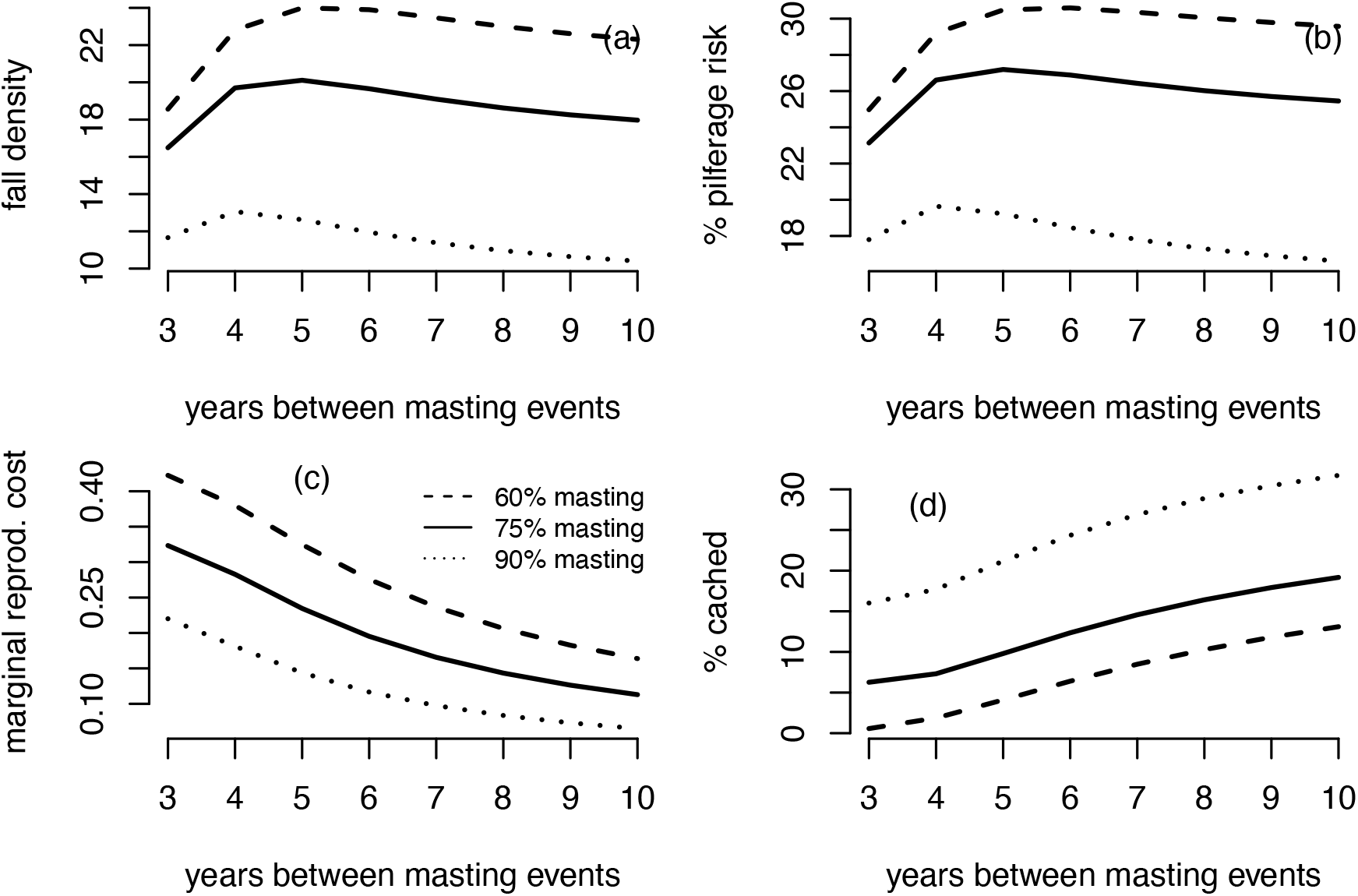
Mast year fall population density (a), pilferage risk (b), marginal reproductive costs of caching (c), and proportion of seeds cached rather than eaten (d) as a function of masting interval. Dashed, solid, and dotted lines represent masting intensity (60, 75 or 90% of total seed production occurring during mast years). All dependent variables are given at the evolutionary stable caching strategy and its associated periodic population dynamics.

**Fig. 4.**
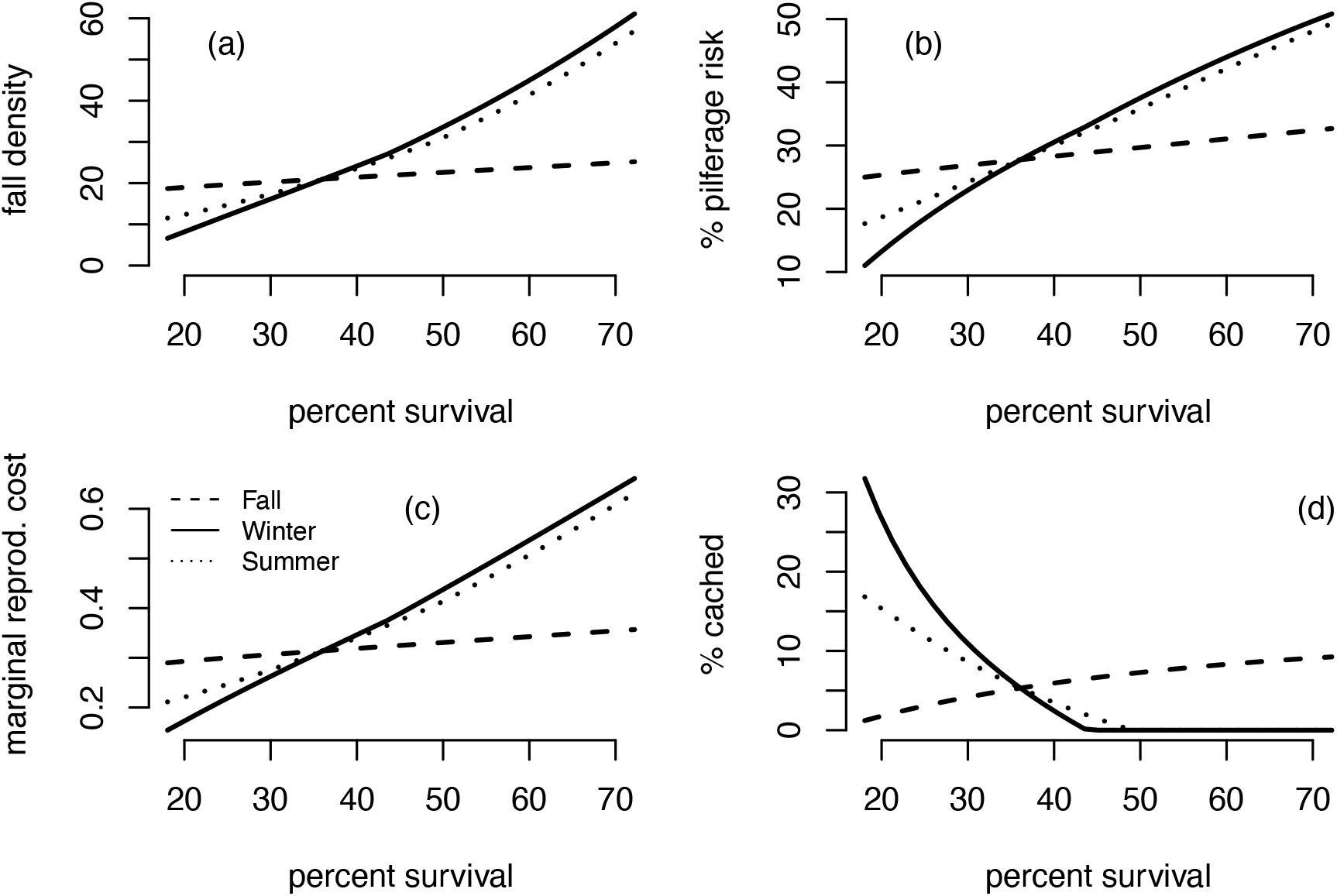
Mast year fall population density (a), pilferage risk (b), marginal reproductive costs of caching (c), and proportion of seeds cached rather than eaten (d) as a function of scatterhoarder survival. Dashed, solid, and dotted lines denote responses to changes in fall, winter/spring, and summer survival, respectively. All dependent variables are given at the evolutionary stable caching strategy and its associated periodic population dynamics.

Furthermore, increasing masting intensity is associated with a decline in marginal reproductive costs of caching – the cost of caching seeds for future use rather than using seeds for current reproduction (Fig. 2c). Lower population densities and higher seed abundance during mast years mean more seeds per individual. Because reproduction is a saturating function of seeds consumed, the marginal reproductive cost of caching declines as seed abundance increases.

As a result of reduced pilferage risk and marginal costs, increasing masting intensity causes an accelerating increase in the ESS proportion of seeds cached rather than eaten (Fig. 2d).

Higher recovery advantage by cache owners reduces pilferage risk (dashed vs. solid vs. dotted lines Fig. 2a), but has little effect on population densities and marginal reproductive costs in the fall of masting years. Consequently, higher owner advantage selects for greater caching.

More years with poor seed crops between masting events lowers marginal reproductive costs (because there are more seeds per individual) (Fig. 3c) but can increase or lower densities of individuals entering the fall of a masting year (Fig. 3a) which increases or lowers the risk of seed pilferage (because more or fewer individuals enter winter) (Fig. 3b). Collectively, the lower reproductive costs outweigh the effects of pilferage risk and select for more caching (Fig. 3d). Varying masting intensity (60, 75 or 90% of seeds produced during mast years: dotted, solid or dashed line on Fig. 3) affects the magnitude of these changes in the manner consistent with Fig. 1., with only minor effects on the shape of responses to the masting interval.

Increasing the survival of scatterhoarders leads to increases in fall population density, pilferage risk, and marginal reproductive costs of caching (Fig. 4). These effects are the strongest due to increases in winter survival, intermediate due to summer survival, and the weakest due to fall survival. This is likely because after masting the greatest concentration of births occurs in the fall and winter, resulting in the winter population having a higher percentage of new individuals (who are not subject to mortality during the previous period) than the summer and fall populations. Thus, an increase in mortality in the fall affects a smaller proportion of the population than an increase in mortality in the winter or summer. Despite increasing marginal reproductive costs and pilferage risk, increasing fall survival, unlike winter/spring or summer survival, selects for more caching. This occurs because, unlike summer or spring/winter survival, fall survival increases the likelihood that an individual caching in the fall will survive to the winter/spring to make use of their cache.

## Discussion

The fact that masting causes strong fluctuations in populations of seed-eating animals has been well-known for a long time (Curran and Leighton 2000; Ostfeld and Keesing 2000; Bogdziewicz et al. 2016), yet the traditional research focus has been on how the satiation-starvation cycle reduces seed losses to pre- and post-dispersal seed predators. More recently, researchers suggested that seed masting is one of the means by which plants manipulate behavior of their dispersers (Vander Wall 2010). According to this reasoning, satiation of current energy needs induces granivores to cache seeds for future use (Vander Wall 2010). Here we show that the effects of masting on population dynamics and caching behavior are mutually dependent. By decreasing the degree of pilfering, the satiation-starvation cycle due to more extreme seed masting events may promote the evolution and maintenance of seed caching behavior. Thus, the decrease in seed predation, increase in per capita scatterhorder satiation, and reduction in pilfering pressure may each represent an important pathway by which the scatterhorder satiation-starvation cycle induced by masting may improve plant recruitment (Fig. 5). These nuanced interactions between plant and seed predator emphasize the importance of studying the feedbacks between population dynamics and behavioral evolution.

**Fig. 5.**
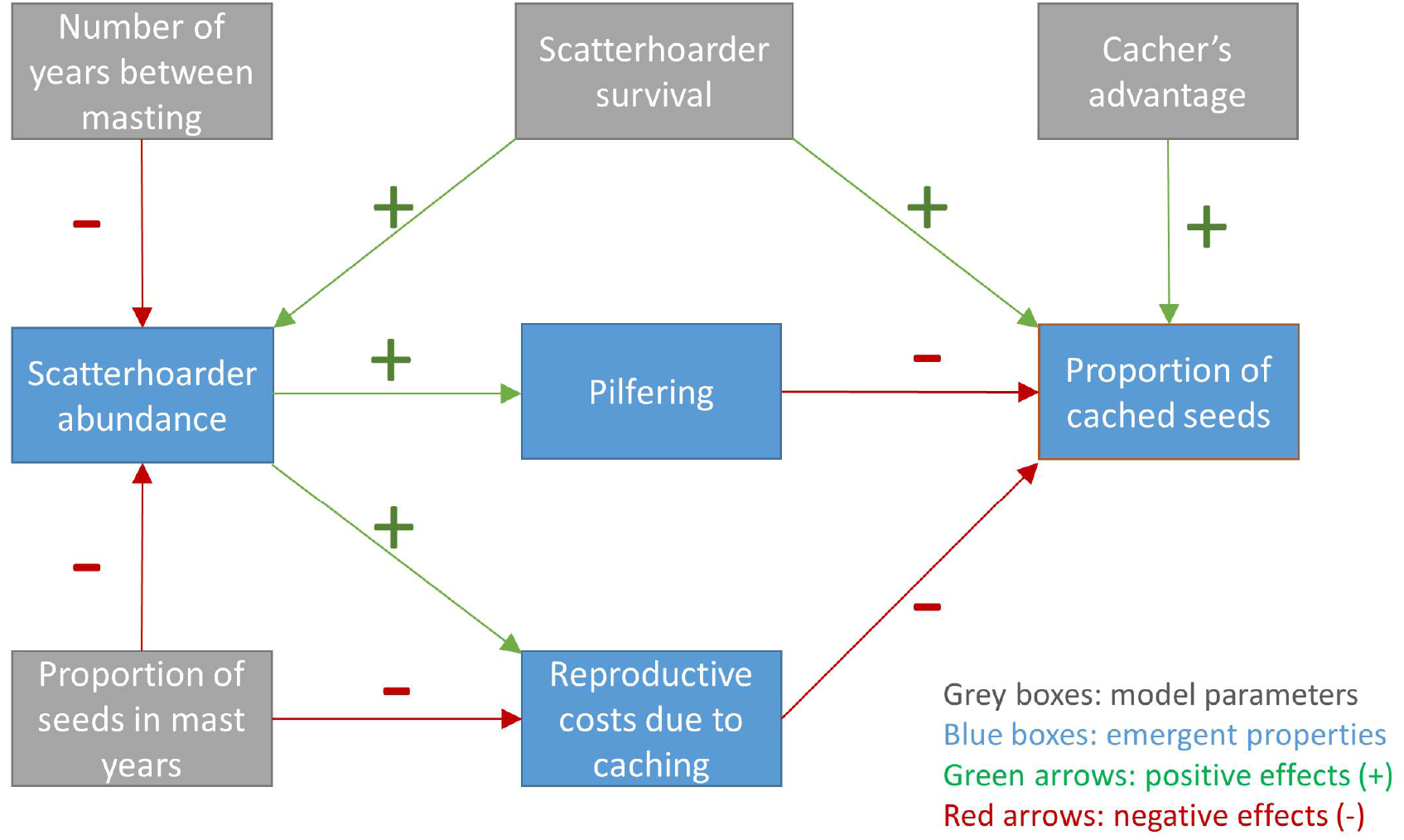
Relationships between input parameters (masting interval and intensity, expressed as the proportion of seeds produced during mast years, scatterhoarder survival, and cacher’s advantage in cache recovery) and emergent properties of the model (scatterhoarder population density, the proportion of cached seeds that are pilfered, reproductive costs of caching, and the quantity of interest: the proportion of seeds cached by scatterhoarders).

Our results suggest that when seed production is highly variable, seed caching can evolve even when cache owners have little advantage over naive foragers in seed recovery (compare with Andersson and Krebs 1978). However, the mechanism that we describe is not mutually exclusive with other evolutionary explanations of scatter-hoarding. It can promote this behavior in synergy with the cache owner’s advantage (Andersson and Krebs 1978) and reciprocal pilferage (Vander Wall and Jenkins 2003).

The costs of cache loss to pilferers are reduced in our model because periods of intense seed production coincide with low densities of scatterhoarders – and thus few potential pilferers (see Dittel and Vander Wall 2018 for experimental data demonstrating that the magnitude of cache pilferage is determined by the abundance of scatterhoarders). When there is pronounced masting with relatively long intervals between masting events, densities of scatterhoarders entering the start of the next large masting event are low (Figs. 2 & 3). Consequently, individuals are able to collect enough seeds to satiate their reproductive needs. As the yearly fitness is determined by the geometric mean of their fitness across the seasons and this geometric mean decreases with variation (Lewontin & Cohen 1969; Gillespie 1977; Schreiber 2015), the benefits of reducing seasonal variation in fitness by increasing winter/spring reproduction (fueled by cached seeds) outweigh the diminishing returns of increasing reproduction in the fall (fueled by immediate seed consumption).

Our results make a prediction that plants dispersed by scatterhoarders should have high interannual variation of seed production (typically measured with coefficient of variation, CV) relatively to plants dispersed by other means. This appears to be the case, at least when plants dispersed by scatterhoarders (synzoochorously) are compared to plants dispersed by frugivores (endozoochorously) (Herrera et al. 1998, Kelly and Sork 2002, Pearse et al. 2020). When explaining this pattern, researchers emphasized contrasting selective pressures acting on these groups of plants. Avoiding the risk of satiating frugivores was suggested as a factor that stabilizes seed production in plants dispersed endozoochorously. On the other hand, variable seed production in synzoochorous plants was interpreted as an adaptation that enabled reducing seed mortality caused by animals that act as seed predators and only incidentally disperse seeds (Herrera et al. 1998). However, we suggest that the high CV of plants dispersed by scatterhoarders can also be linked to the caching behavior of scatterhoarders (see also Lichti et al. 2020 for a model exploring the connection between caching behavior and seed trait evolution).

If, as our simulations suggest, masting intensity and mast interval are important for seed caching, then changes in plant masting patterns might affect the dynamics of seed caching, and therefore also the recruitment in plant populations. Our model is loosely based on the European beech – *Apodemus* mice system (Jensen 1982; Zwolak et al. 2016). Several studies have suggested that the European beech shows more frequent masting in recent years, probably due to global warming (Kantorowicz 2000; Schmidt 2006; Övergaard et al. 2007; Bogdziewicz et al. 2020). This could shift the beech-rodent interactions towards antagonism, with higher rodent abundances (predicted also by Imholt et al. 2013), more seed consumed and fewer cached (recall that caching declines with more frequent masting: Fig. 3). On the other hand, a recent meta-analysis of global data suggests that masting has become more pronounced (Pearse et al. 2017). Such a change could make seed caching more profitable for granivores (higher intensity of masting promotes caching: Fig. 2). However, extreme interannual variation in seed crops might lead to a decline and even extinction in granivore populations, due to the difficulty in tracking resource levels.

Moreover, any environmental change that affects scatterhoarder population dynamics could alter caching behavior and, thereby, impact seed mortality. For example, we found that increased scatterhoarder survivorship during the winter or summer may select against caching behavior by increasing population densities entering the masting years (Fig. 4). Thus, changes in winter or summer conditions that are favorable for mice could harm seedling recruitment both directly by increasing seed predation and indirectly by discouraging seed caching. In contrast, autumn conditions that are favorable for mice are likely to improve seedling recruitment because increased fall survivorship of scatterhoarders selects for more caching.

Just like every model, the one presented here simplifies reality. For example, in many ecosystems different masting species co-occur. Such species often mast synchronously due to shared climatic drivers (Curran et al. 1999; Kelly and Sork 2002; Schauber et al. 2002; Shibata et al. 2002; Bogdziewicz et al. 2018). If seed production is synchronous, the consequences for scatterhoarders will be similar to masting by one tree species. However, if masting is asynchronous, its outcome might be similar to reducing masting interval (Fig. 3d), i.e. the selective pressure to cache seeds will be weaker.

Furthermore, populations of scatterhoarders that have relatively high survival and low reproduction (e.g. corvids) might not fluctuate in response to masting as strongly as do populations of more productive scatterhoarders, such as chipmunks (Bergeron et al. 2011), squirrels (McShea 2000; Selonen et al. 2015), or mice (Pucek et al. 1993, Wolff 1996; Falls et al. 2007). However, even in species such as corvids, masting can affect the benefits of caching through similar mechanisms, i.e. reduced risk of interspecific seed pilferage (due to decreased abundance of sympatric rodents: e.g. Thayer and Vander Wall 2005) and lower marginal reproductive costs of caching.

Examining ultimate causes and ecological determinants of caching behavior will help to understand former selective pressures on synzoochorous plants, current dynamics of seed dispersal, and future alterations in seed dispersal patterns caused by global changes. Our study provides a step in this direction and suggests several promising avenues for prospective research. For example, future work should address the evolution of caching reaction norms instead of the simple threshold for caching considered here. Additionally, evolution of caching strategies could be different when individual variation in personalities or, more generally, phenotypes of seed dispersing animals (Zwolak 2018, Zwolak and Sih 2020) is taken into account. Finally, future studies could examine these interaction from the plant perspective, for example by determining masting patterns that maximize seedling recruitment.

## Data accessibility

The code for the main functions of the model, model results, and sensitivity analyses is included as Supplementary Data.

## Authors’ contributions

RZ and SS conceived the study. SS and DC developed and analyzed the model with feedback from RZ and AS. RZ wrote the first draft of the manuscript. All authors critically revised the draft and approved the final version of the article.

## Acknowledgments

Our manuscript benefitted from comments by Dave Kelly and two anonymous reviewers, and discussions with Michał Bogdziewicz.

## Appendix A

**Table of Mathematical Notation**

**Table A1.**
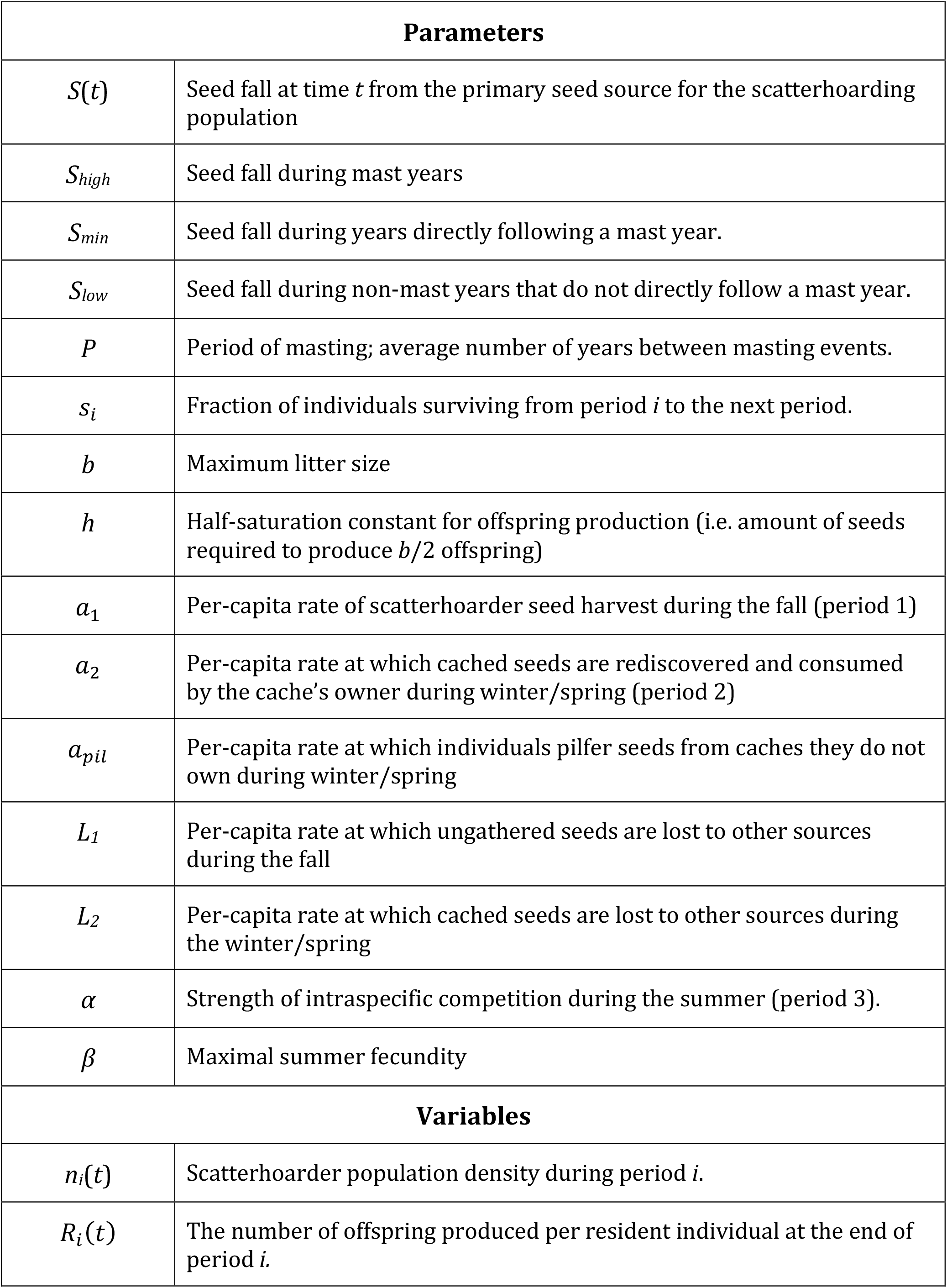
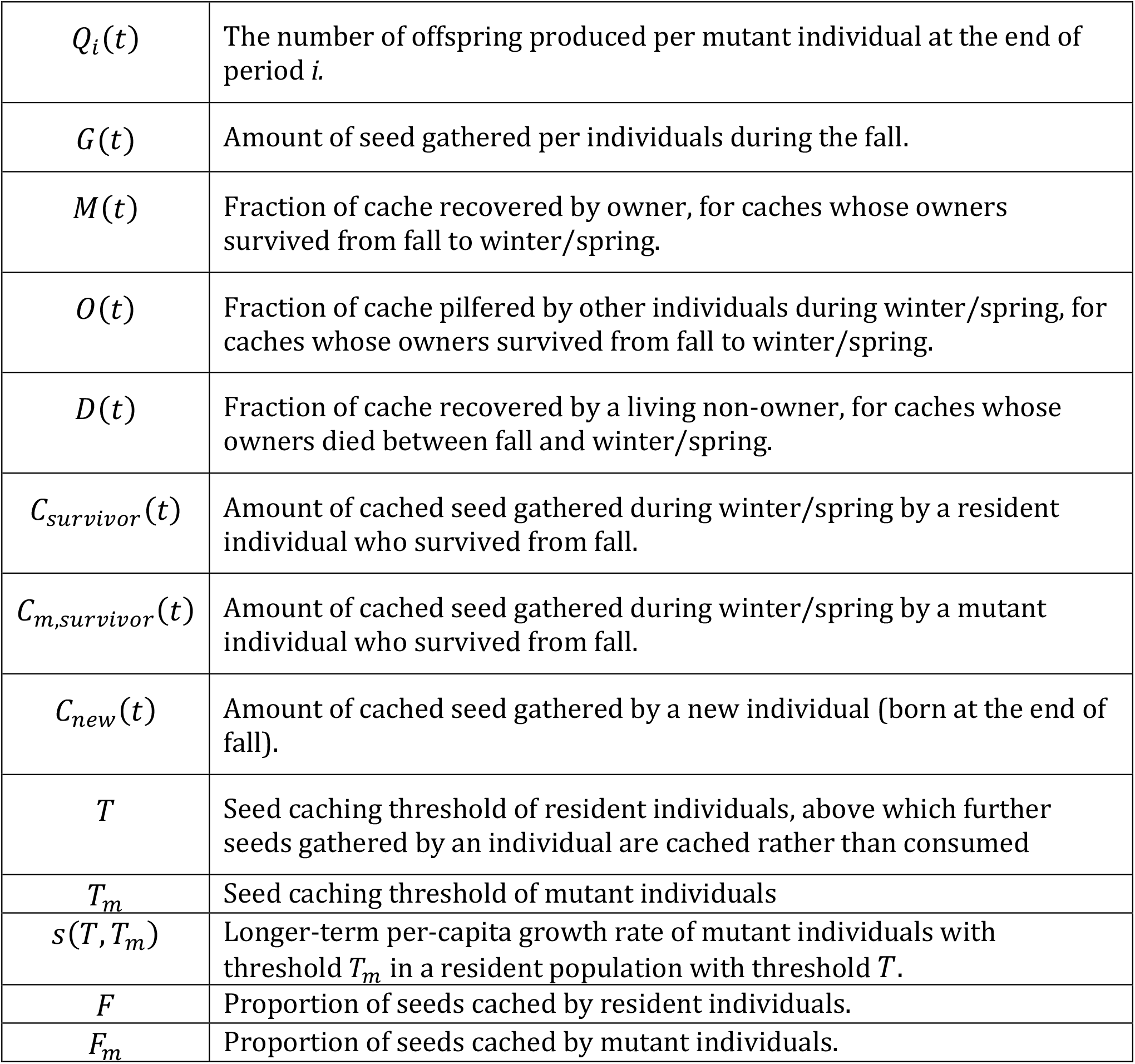
A list of names and descriptions for the parameters and variables introduced in the main text.

| **Parameters** |  |
| --- | --- |
| $S(t)$ | Seed fall at time $t$ from the primary seed source for the scatterhoarding population |
| $S_{high}$ | Seed fall during mast years |
| $S_{min}$ | Seed fall during years directly following a mast year. |
| $S_{low}$ | Seed fall during non-mast years that do not directly follow a mast year. |
| $P$ | Period of masting; average number of years between masting events. |
| $s_i$ | Fraction of individuals surviving from period $i$ to the next period. |
| $b$ | Maximum litter size |
| $h$ | Half-saturation constant for offspring production (i.e. amount of seeds required to produce $b/2$ offspring) |
| $a_1$ | Per-capita rate of scatterhoarder seed harvest during the fall (period 1) |
| $a_2$ | Per-capita rate at which cached seeds are rediscovered and consumed by the cache's owner during winter/spring (period 2) |
| $a_{pil}$ | Per-capita rate at which individuals pilfer seeds from caches they do not own during winter/spring |
| $L_1$ | Per-capita rate at which ungathered seeds are lost to other sources during the fall |
| $L_2$ | Per-capita rate at which cached seeds are lost to other sources during the winter/spring |
| $\alpha$ | Strength of intraspecific competition during the summer (period 3). |
| $\beta$ | Maximal summer fecundity |
| **Variables** |  |
| $n_i(t)$ | Scatterhoarder population density during period $i$. |
| $R_i(t)$ | The number of offspring produced per resident individual at the end of period $i$. |
| $Q_i(t)$ | The number of offspring produced per mutant individual at the end of period $i$. |
| $G(t)$ | Amount of seed gathered per individuals during the fall. |
| $M(t)$ | Fraction of cache recovered by owner, for caches whose owners survived from fall to winter/spring. |
| $O(t)$ | Fraction of cache pilfered by other individuals during winter/spring, for caches whose owners survived from fall to winter/spring. |
| $D(t)$ | Fraction of cache recovered by a living non-owner, for caches whose owners died between fall and winter/spring. |
| $C_{survivor}(t)$ | Amount of cached seed gathered during winter/spring by a resident individual who survived from fall. |
| $C_{m,survivor}(t)$ | Amount of cached seed gathered during winter/spring by a mutant individual who survived from fall. |
| $C_{new}(t)$ | Amount of cached seed gathered by a new individual (born at the end of fall). |
| $T$ | Seed caching threshold of resident individuals, above which further seeds gathered by an individual are cached rather than consumed |
| $T_m$ | Seed caching threshold of mutant individuals |
| $s(T, T_m)$ | Longer-term per-capita growth rate of mutant individuals with threshold $T_m$ in a resident population with threshold $T$. |
| $F$ | Proportion of seeds cached by resident individuals. |
| $F_m$ | Proportion of seeds cached by mutant individuals. |

## Appendix B

**Global Sensitivity Analysis**

We explored the robustness of results in Figures 2, 3, and 4 of the main text to base-line parameter choices. For each Figure, we reran the simulations for 100 randomly sampled values of the parameters (except the manipulated parameter on the horizontal axis). Parameters, say with a default value of x, were randomly sampled independent of one another from a uniform distribution on the interval [x/1.5,1.5x]. For Figure 4, we only varied the fall survival parameter on the horizontal axis, as only this survival parameter gave a more nuanced result in the main text. Each of these sensitivity analyses confirmed the general conclusions for the main text. Increasing masting intensity led to lower fall densities, pilferage risk, marginal reproductive costs, and more caching during the masting year (Fig. B1). Increasing the number of years between masting events lead to lower marginal reproductive costs and more caching during the masting year (Fig. B2). In contrast, increasing fall survival led to more caching despite higher fall densities, pilferage risk, and marginal reproductive costs during the masting year (Fig. B3).

**Fig. B1.**
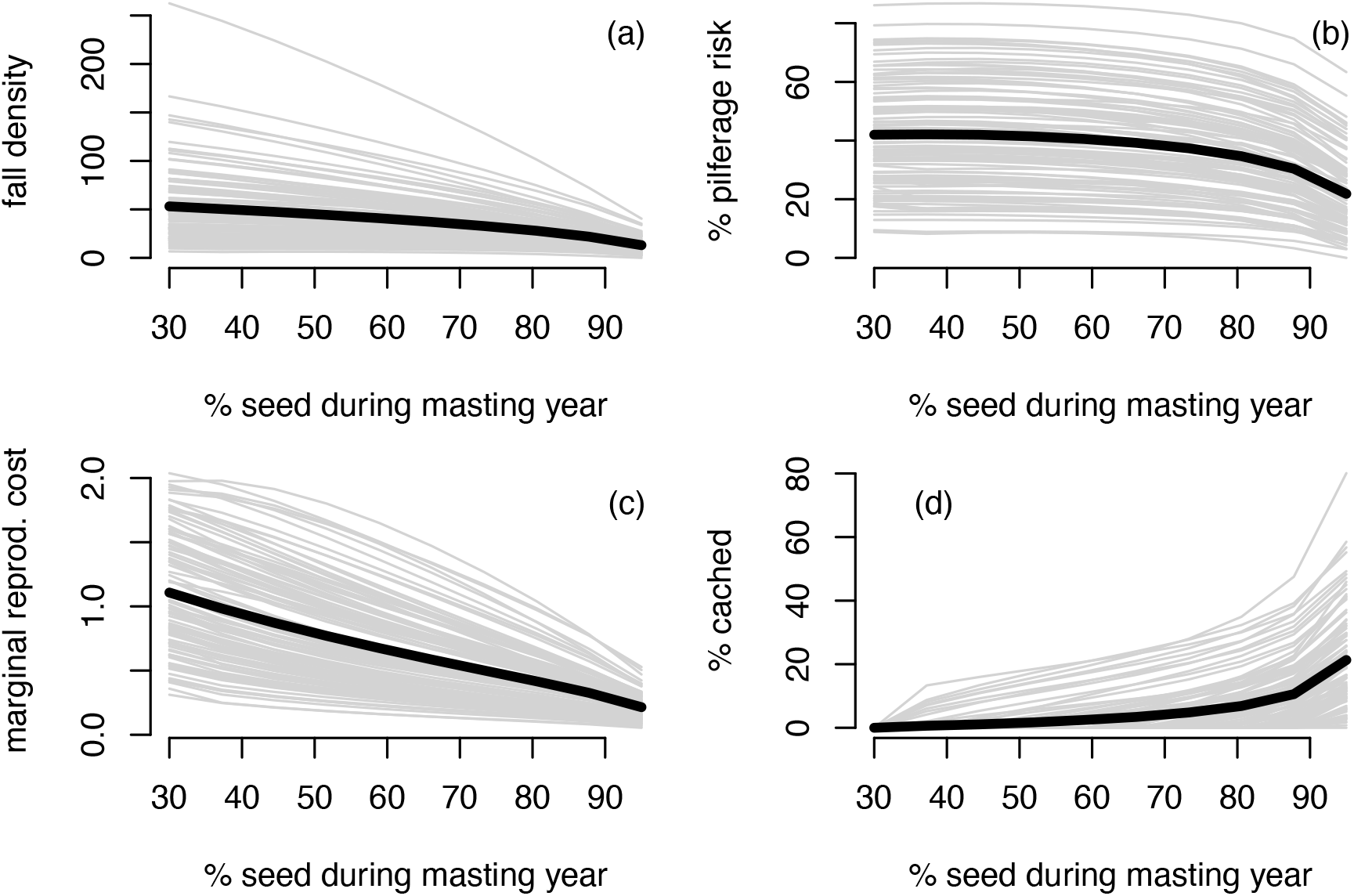
Mast year fall population density in individuals/ha (a), pilferage risk, defined as the probability that cached seed would be pilfered (b), marginal reproductive costs of caching (c), and proportion of seeds cached rather than eaten (d) as a function of masting intensity (expressed as the percentage of total seed production that occurs during mast years), with mast occurring every 4^th^ year.

**Fig. B2.**
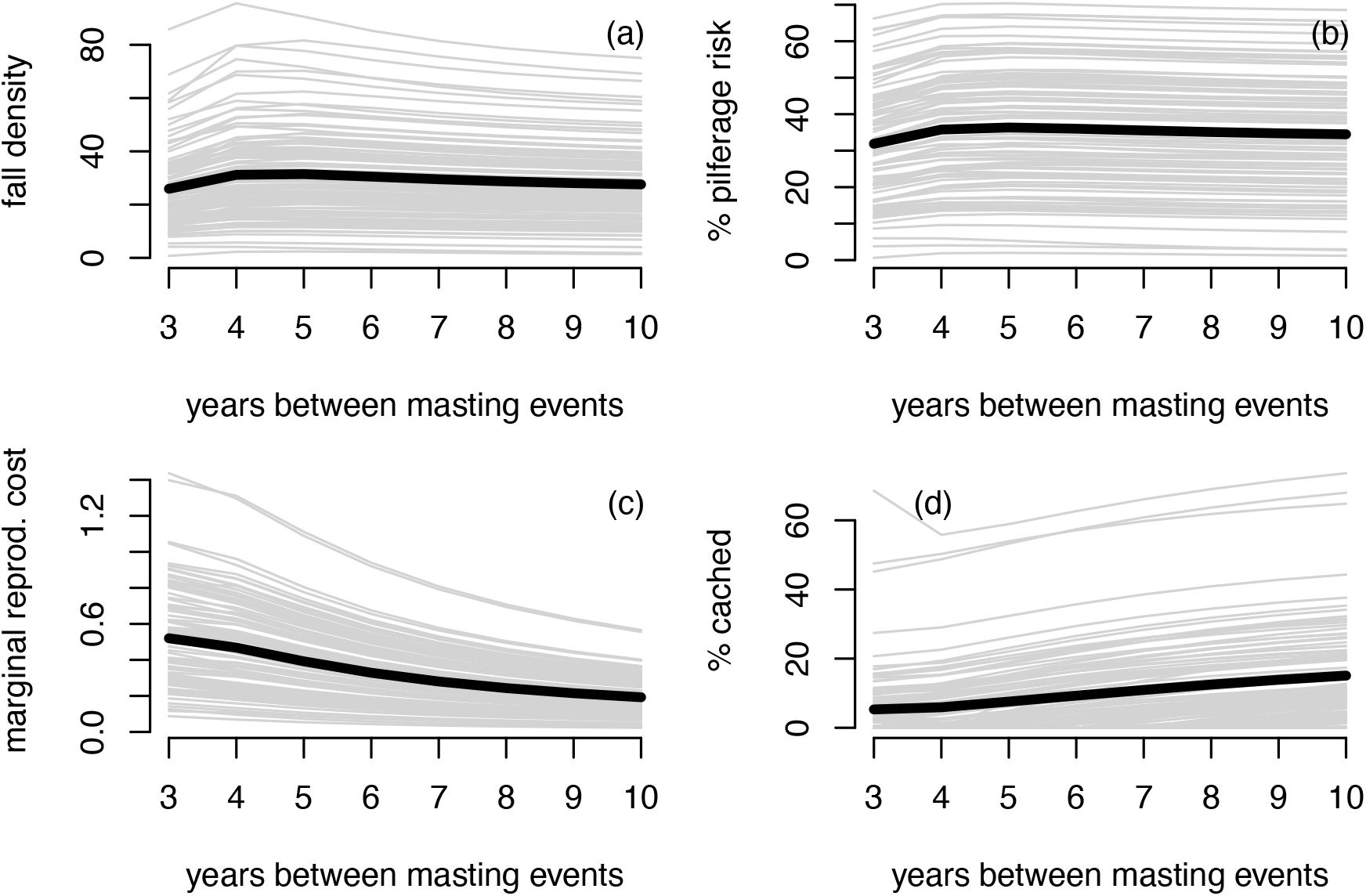
Mast year fall population density (a), pilferage risk (b), marginal reproductive costs of caching (c), and proportion of seeds cached rather than eaten (d) as a function of masting interval. Plotted variables represent values in the mast year.

**Fig. B3.**
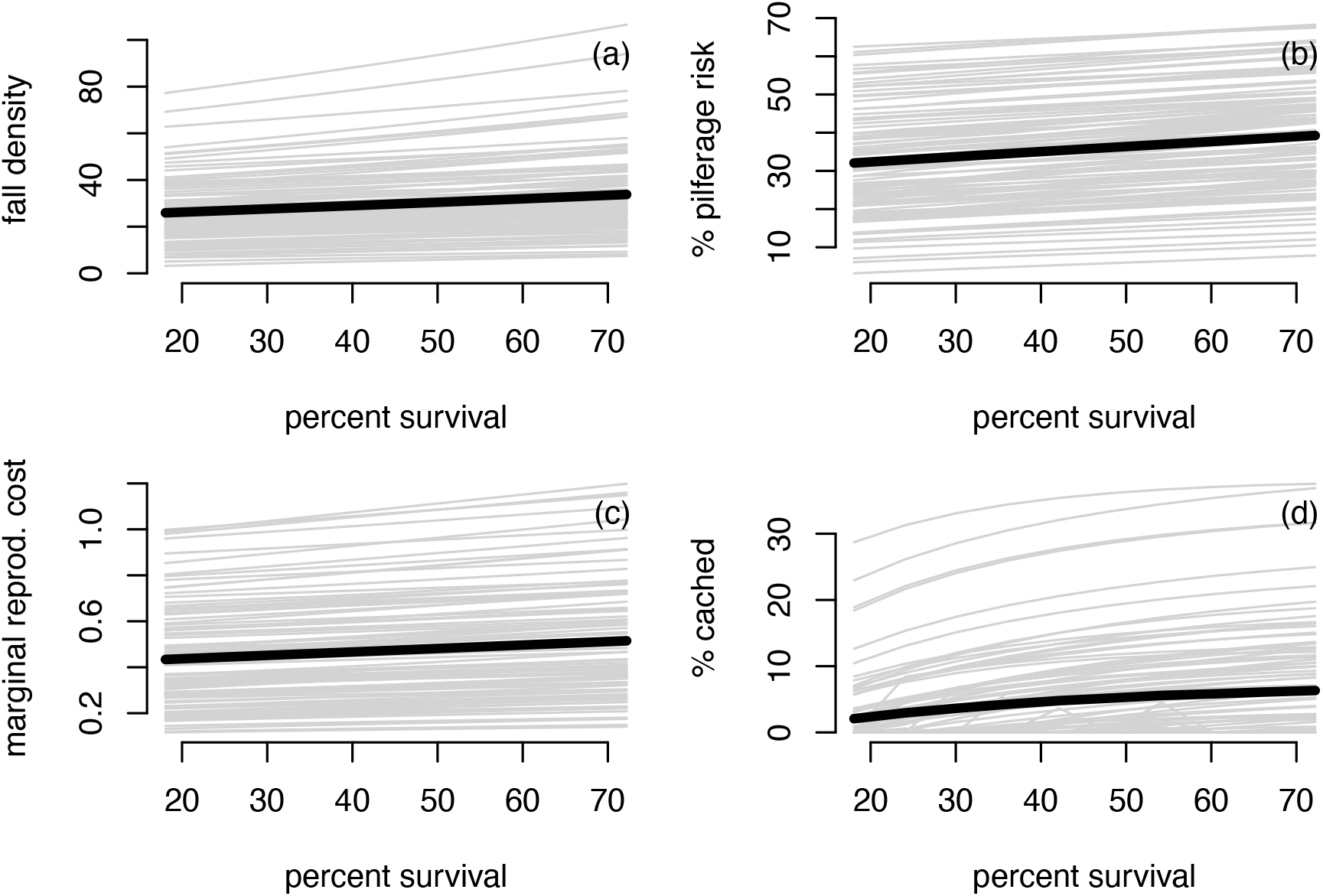
Mast year fall population density (a), pilferage risk (b), marginal reproductive costs of caching (c), and proportion of seeds cached rather than eaten (d) as a function of scatterhoarder survival in the fall.

## Appendix C

**Derivation of the fitness gradient**

To derive the fitness gradient, assume that the population dynamics of individuals with the resident threshold strategy *T* converges to a periodic solution of period *kP* : *n*_*i*_(1), *n*_*i*_(2), …, *n*_*i*_(*p*) for *i* = 1, 2, 3. The more general case of aperiodic dynamics is discussed at the end of this appendix. For each 1 *≤ t ≤ kP*, let *G*(*t*) be the amount seeds gathered in Fall by each resident individual in year *t*, and *Q*_1_(*t*), *Q*_2_(*t*), *Q*_3_(*t*) be the number of offspring produced by an individual playing the mutant strategy *T*_*m*_ during the Fall, Winter/Spring, and Summer, respectively, in year *t*. Then, the long-term per-capita growth rate of the mutant is

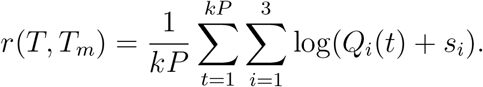

Taking the derivative with respect to *T*_*m*_ and evaluating at *T*_*m*_ = *T* gives us the fitness gradient:

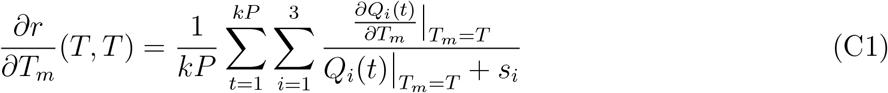

As 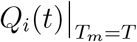 corresponds to the fecundity of resident individuals,

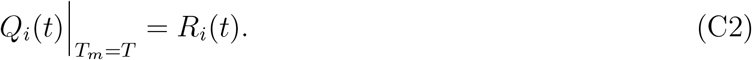

It remains to compute the partial derivatives of the *Q*_*i*_(*t*) terms. For *i* = 1, we have (from the main text)

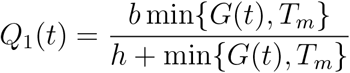

is piecewise defined depending on whether *G*(*t*) *< T* or *> T*. Specifically, we get

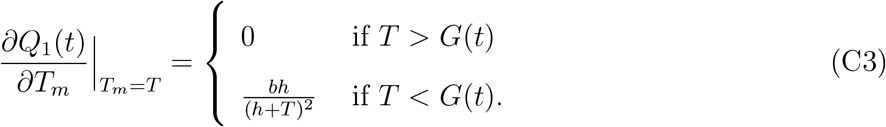

For *i* = 2, we have (from the main text)

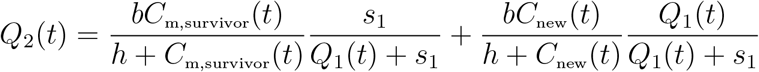

where *Q*_2_(*t*) differs from *R*_2_(*t*) only in its first term due to surviving individuals with the mutant caching strategy:

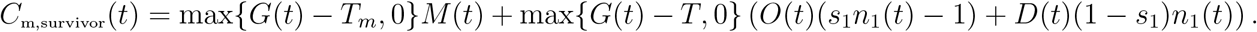

Hence, by chain rule,

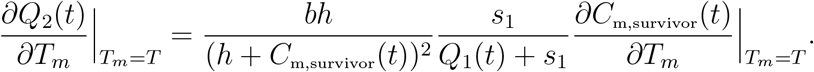

At *T*_*m*_ = *T*, we have *C*m,survivor(*t*) = *C*survivor(*t*) and *Q*_1_(*t*) = *R*_1_(*t*). Furthermore,

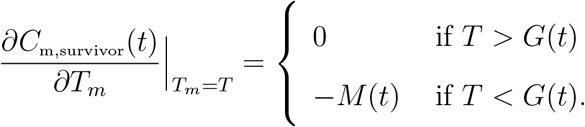

Thus,

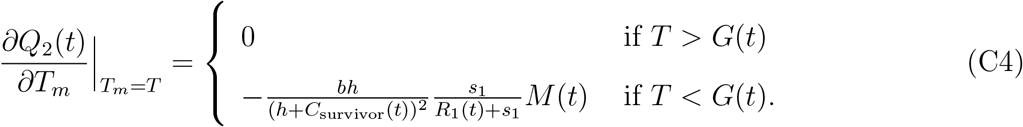

Finally, as discussed in the main text, *Q*_3_(*t*) = *R*_3_(*t*) and, consequently, doesn’t depend on *T*_*m*_. Hence,

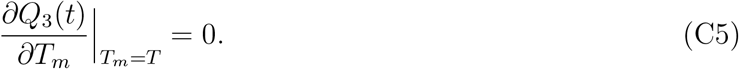

Substituting equations (C2)–(C5) into (C1) provides an explicit expression for the fitness gradient. These calculations also apply to the more general case when the resident dynamics are asymptotically stationary but not necessarily periodic. By asymptotic stationarity, we mean that there exists a probability measure *µ* on the (*S, n*_1_) state space such that

**Figure C1:**
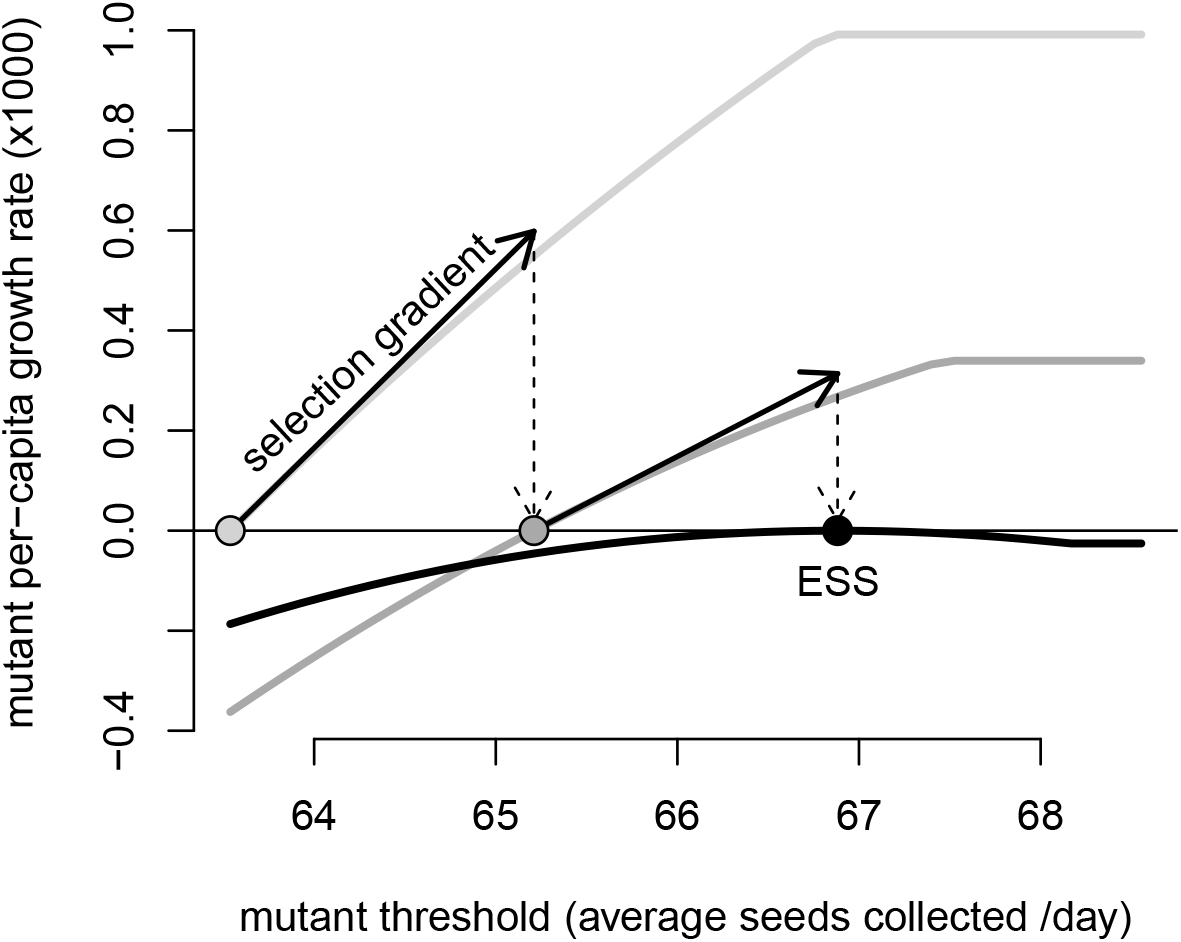
Convergence to the evolutionary stable strategy (ESS) by following the selection gradient. The per-capita growth rates r(*T, T*_*m*_) of mutants playing threshold strategy *T*_*m*_ (measured in average # seeds collected per day) plotted (light gray to black curves) for different resident threshold strategies *T* (light gray to black circles) including the ESS strategy (black circle). The selection gradient corresponds to the slope of the arrows. When the resident strategy corresponds to the ESS (black circle), the per-capita growth rate of the mutant is maximized at the ESS value. Parameter values as in Figure 1 from main text.

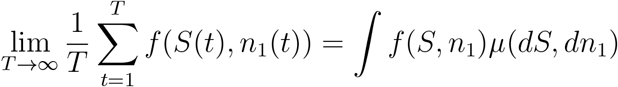

for all continuous functions *f* (*S, n*_1_) i.e. the Birkhoff averages converge. In this case, the per-capita growth rate of the mutant is well-defined and equals

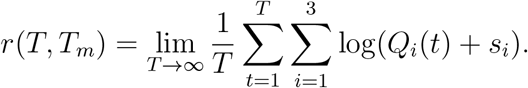

The selection gradient equals

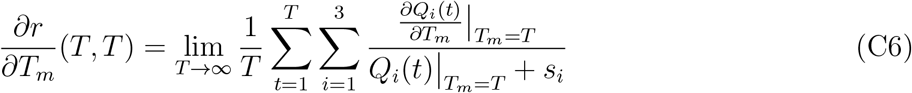

and can be explicitly evaluated by using (C2)–(C5). Figure C1 illustrates how the selection gradient can be used to find the ESS.

